# Characterizing Traction Forces in Upstream-Migrating Hematopoietic-like KG1a Cells Under Shear Flow

**DOI:** 10.1101/2025.06.13.659511

**Authors:** Dong-hun Lee, Daniel A. Hammer

## Abstract

Cell migration is critical to leukocyte function, enabling leukocytes to patrol tissues and respond to inflammatory cues. Upstream migration is a distinct form of cell motility which enables leukocytes to move against the direction of fluid flow on Intercellular Adhesion Molecule-1 (ICAM-1) surfaces. Upstream migration is mediated by the leukocyte integrin, Lymphocyte Function-Associated Antigen-1 (LFA-1). While this behavior has been observed across multiple immune cell types, the mechanical forces underlying upstream migration have not been measured. Here, we demonstrate the use of Traction Force Microscopy (TFM) to quantify spatiotemporal patterns of force generation during upstream migration of KG1a cells, a hematopoietic progenitor cell line that exhibits robust upstream migration on ICAM-1 functionalized hydrogels.

Under static (no-flow) conditions, KG1a cells displayed random motility and traction profiles that varied with time. In contrast, cells exposed to shear flow generated persistent, polarized tractions aligned with the axis of migration. Population analysis showed that maximum RMS traction forces were significantly elevated during upstream migration compared to static conditions (mean: 428.5 ± 63.0 nN vs 220.8 ± 22.2 nN, p = 0.0078), as were average RMS forces (mean: 82.6 ± 12.9 nN vs 45.9 ± 4.4 nN, p = 0.0184), while minimum force values remained comparable between groups. These findings indicate a specific amplification of stresses required to overcome applied forces during upstream migration.

By integrating single-cell and population-level force analyses, this study defines upstream migration as a mechanically reinforced state characterized by amplified, directionally coherent traction dynamics. Our methods enable future dissection of the molecular regulators that coordinate force generation with migration under flow.

**SIGNIFICANCE:** Upstream migration is a critical yet poorly understood mode of immune cell motility. Here, we use traction force microscopy under physiologically relevant shear stresses to provide the first quantitative characterization of traction dynamics during upstream migration. We show that upstream migrating cells generate significantly greater average and peak forces than cells under static conditions, revealing a distinct mechanical program for directed migration against flow. These findings establish a platform for dissecting the molecular regulators of force generation in immune cells and set the stage for future perturbation-based studies aimed at understanding how mechanical forces shape immune cell trafficking and vascular navigation.

## INTRODUCTION

Cell migration is a vital process that underpins numerous physiological and pathological phenomena, such as immune surveillance and cancer metastasis (1). Leukocytes, as key players in immune function, exhibit remarkable, diverse motility, allowing them to migrate efficiently through vascular environments and infiltrate tissues to combat infections and maintain homeostasis (2). Tissue entry is orchestrated through a well-characterized sequence known as the leukocyte adhesion cascade, which progresses through the steps of capture, rolling, migration, and transmigration (3, 4). Each step involves distinct molecular and mechanical cues that guide leukocytes to their destination (5, 6). Of particular interest in this paper is the migration phase, where cells exhibit diverse patterns of motility affected by both biochemical signals and mechanical stimuli (7, 8).

A striking example of mechanosensitive migration is upstream migration — cellular movement against the direction of external shear flow. First identified in T cells by Valignat and coworkers in 2013 (9), upstream migration highlights the interplay between integrin signaling and external forces. This phenomenon has been shown to be mediated by the interaction between the integrin Lymphocyte Function-Associated Antigen-1 (LFA-1) and its ligand Intercellular Adhesion Molecule-1 (ICAM-1), while the Very Late Antigen-4 (VLA-4)/Vascular Cell Adhesion Molecule-1 pair favors downstream migration (9–11). Similar behavior has been observed in Hematopoietic Stem and Progenitor cells (HPSCs) and an immortalized human surrogate cell line, the KG1a cells. Upstream migration was also observed in neutrophils, and the neutrophil-like cell line, HL-60 cells, when Mac-1 integrin was blocked (12–15). Despite the recurring observation of upstream migration across different leukocyte types, the forces involved in this behavior have not been quantified.

Previous studies investigating upstream migration have primarily relied on qualitative or semi-quantitative metrics, such as the fraction of cells migrating upstream, the formation of lamellipodia or uropodia, or overall directional persistence. For instance, Roy et al. investigated the role of LFA-1 and the Crk adaptor proteins using actin staining and imaging-based directional indices to infer cytoskeletal polarization and migratory behavior under flow (11, 16). While these studies reveal important aspects of upstream motility, they do not offer direct measurements of the forces leukocytes exert during upstream migration. As a result, it remains rather unclear whether upstream migration involves greater mechanical output than migration in the absence of flow, or how traction forces are distributed spatially and temporally during this behavior. Quantitative assessment of these forces is therefore needed to establish a clear baseline for future mechanistic studies.

Traction Force Microscopy (TFM) provides a robust platform for addressing this gap. By embedding fluorescent beads within polyacrylamide substrates and tracking bead displacements over time, TFM enables high-resolution mapping of the forces cells exert on their substrate. (17). Quantitative outputs include not only spatially resolved force vectors, but also global measures such as root-mean-square (RMS) force. TFM has proven particularly valuable in analyzing motility in various immune cells, including during neutrophil chemotaxis and macrophage polarization (18–20). However, TFM has not been used to study traction generation during upstream migration under shear flow, a key omission given that upstream migration likely involves distinct mechanical demands in comparison to other modes of trafficking. The ability to extract force values from adherent, motile, shear-responsive cells would represent a technical advance.

In this study, we apply TFM to quantify the traction forces generated by KG1a cells during both static (no-flow) conditions and upstream migration against shear flow on ICAM-1 functionalized polyacrylamide hydrogels. KG1a cells are particularly well suited for this study because they exhibit robust upstream migration, form transient yet measurable adhesions on hydrogels, and offer experimental consistency compared to primary cells (14). In addition, they are a well-documented surrogate for HPSCs (14), and because they are grow in culture, they can be easily manipulated through knockdowns and gene editing. We employed a well-defined shear rate of 400 s^−1^, which lies within the physiological range observed in postcapillary venules (11), to induce upstream migration and analyzed population-level force distributions across both conditions. Unlike previous studies that inferred mechanical engagement indirectly through morphology or directionality, we report direct measurements of minimum, maximum, and average traction forces in each condition. Our findings reveal a clear increase in average and peak forces during upstream migration, suggesting that enhanced mechanical output is a defining feature of this behavior. Together, these results establish a quantitative benchmark for future studies and offer a foundation for investigating how force-sensitive pathways regulate migration under shear flow.

## MATERIALS AND METHODS

### KG1a Cell Culture

KG1a cells (ATCC, Manassas, VA) were cultured in 25 cm^2^ culture flasks with Iscove’s Modified Dulbecco’s Medium (IMDM; Invitrogen, Carlsbad, CA) supplemented with 20% Fetal Bovine Serum (FBS; Corning, Corning, NY). Cells were maintained at 37°C with 5% CO_2_ in a humidified incubator.

To harvest cells, the cultures were centrifuged at 125 x g for 7 minutes. The pellet was resuspended in running media composed of IMDM supplemented with 1% D-glucose (Sigma-Aldrich, St. Louis, MO) and 0.5% Bovine Serum Albumin (BSA; Sigma-Aldrich). Harvested cells were used immediately for experiments to maintain viability and consistency across experiments.

### Preparation of Hydrogel Substrates

10 kPa Polyacrylamide hydrogels were prepared as described previously by Jannat and coworkers, with slight modifications (20). Precleaned glass slides (25 × 75mm; Thermo Fisher) were soaked in 0.1M NaOH solution and were left to air dry. A 200 *µ*l aliquot of 3-aminopropyltrimethoxysilane (Sigma-Aldrich, St. Louis, MO) was evenly spread on the glass slide and incubated for 5 minutes. Slides were then washed three times with distilled water and incubated for 30 minutes in 0.5% glutaraldehyde (Sigma-Aldrich) in PBS (Invitrogen). They were then washed again three times with distilled water.

Gels were cast with a final composition of 7.5% acrylamide and 0.15% bis-acrylamide, using 35 mM HEPES buffer (Invitrogen), to produce a Young’s modulus of 10 kPa as reported in Yeung et al. (21). Polymerization was initiated by adding 0.1% w/v TEMED (Bio-Rad) and ammonium persulfate (APS; Bio-Rad). Gel solution was pipetted onto the functionalized slide and overlaid with a Rain-X coated coverslip to create a flat surface. After polymerization, gels were gently detached under water and rinsed thoroughly.

### Gel Functionalization

To functionalize gels, N6 cross-linker was prepared following the protocols outlined by Pless et al (22). and added to gel solutions prior to polymerization. Functionalization steps were adapted from Kim and Hammer (23). Polyacrylamide gels with N6 linkers in it were casted onto a silanized glass slide and were allowed to set using 0.1% w/v ammonium persulfate and TEMED. After polymerization and rinsing, gels were incubated with 10 *µ*g/ml protein A/G (BioVision, Milpitas, CA) for 2 hours at room temperature. Unreacted N6 was blocked by incubating gels with ethanolamine (1/100 parts) in 50 mM HEPES buffer for 30 minutes. Finally, gels were functionalized with 10 *µ*g/ml of ICAM-1 Fc chimeras (R&D Systems, Minneapolis, MN) either for 2 hours at room temperature or for overnight at 4°C.

### Cell Tracking and Quantification of Motility

Images of cells and hydrogels were acquired using a Nikon TE 300 microscope equipped with a Nikon 10x, numerical aperture (NA) 0.25 objective for surface verification. During imaging, cells were maintained at 37°C and 5% CO_2_ using a stage-top incubator. Time-lapse images were acquired at 1-minute intervals for a total of 30 minutes.

Individual cell trajectories were generated using the Manual Tracking plugin in ImageJ (NIH, Bethesda, MD). These trajectories were processed in MATLAB (MathWorks, Natick, MA) to compute migration indices, speed and other motion statistics. The Migration Index (MI) was calculated as the ratio of net displacement along the x-axis to the total path length. MI values range from -1 (perfect upstream migration) to +1 (perfect downstream migration), with 0 indicating unbiased or random motility.

### Traction Force Microscopy

TFM was performed using a Nikon TE 300 inverted microscope with a 40x, 0.45 NA objective, following protocols by Jannat et al. (20), Smith et al. (18), and Reinhart-King et al. (24). Red fluorescent beads (1 *µ*m FluoSpheres, Invitrogen; excitation/emission 580/605 nm) were embedded in the gel surface during polymerization. Time-lapse images of phase contrast and fluorescent channels were acquired every minute for 30 minutes. At the end of each experiment, the field was flushed with running media to remove the cells, and a final image of the relaxed (unstressed) gel was captured.

Shear flow was applied using a motor-powered syringe pump with a built-in control interface, allowing the pump to maintain constant output throughout the experiment. A shear rate of 400 s^−1^ was applied, which lies within the physiological range observed in postcapillary venules (11). Flow calibration and validation followed methods described in Kim and Hammer (23). Experiments were conducted using a custom-built flow chamber designed in-house to support live-cell imaging under flow. (Figure **??**)

Bead displacements were analyzed using custom-written LIBTRC software. Traction stresses were reconstructed using the algorithm described by Dembo and Wang (25). The root-mean-squared (RMS) traction force was computed by integrating the magnitude of traction vectors across the cell footprint:

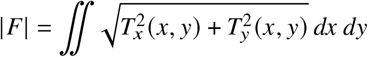

where *T*(*x, y*) = [*T*_*x*_(*x, y*), *T*_*y*_(*x, y*)] represents the continuous field of traction vectors defined at any spatial position *x, y* within the contact region. Population-level distributions of minimum, maximum, and average force values were obtained across static and upstream migration conditions.

## RESULTS

### KG1a cells migrate upstream under shear flow on ICAM-1 functionalized hydrogels

To validate that our ICAM-1 functionalized polyacrylamide substrates support upstream migration, we introduced KG1a cells into a custom-built microfluidic chamber designed for live-cell imaging under flow (Figure **??**). The cells adhered to the hydrogel surface and were exposed to controlled shear flow generated by a syringe pump. Under static conditions, KG1a cells displayed random migration patterns with no discernible preference in directionality. We tracked individual cells over a 30-minute period using time-lapse imaging. As illustrated in Figure 2a, the observed trajectories were consistent with random motility. Quantitative analysis of the MI showed a mean of -0.007 ± 0.053 under static conditions (Figure 2c). A representative video of random migration is available in the supplement (Video S1).

**Figure 1:**
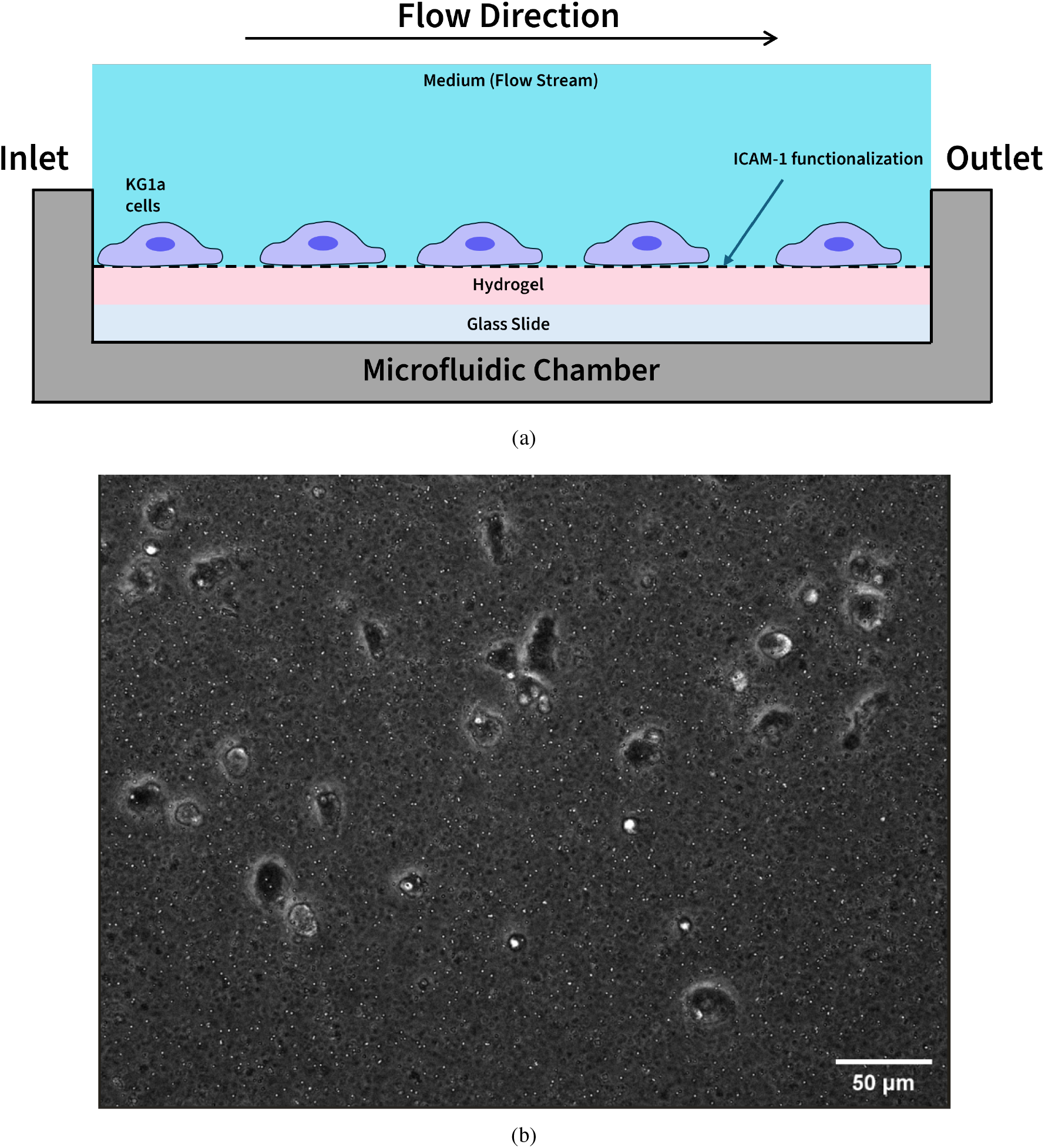
Schematic of the flow chamber setup and sample microscope image. (A) KG1a cells were seeded onto ICAM-1 functionalized polyacrylamide hydrogels within a custom-built microfluidic chamber mounted on a glass slide. Cells adhered to the gel and were subjected to controlled flow at 400 s^−1^ via perfused medium. The flow direction is indicated by the arrow. (B) Representative phase contrast image of KG1a cells adhered to the hydrogel surface under flow. Cells appear spread and partially elongated, consistent with transient adhesion and polarization during upstream migration. Scale bar: 50 *µ*m.

**Figure 2:**
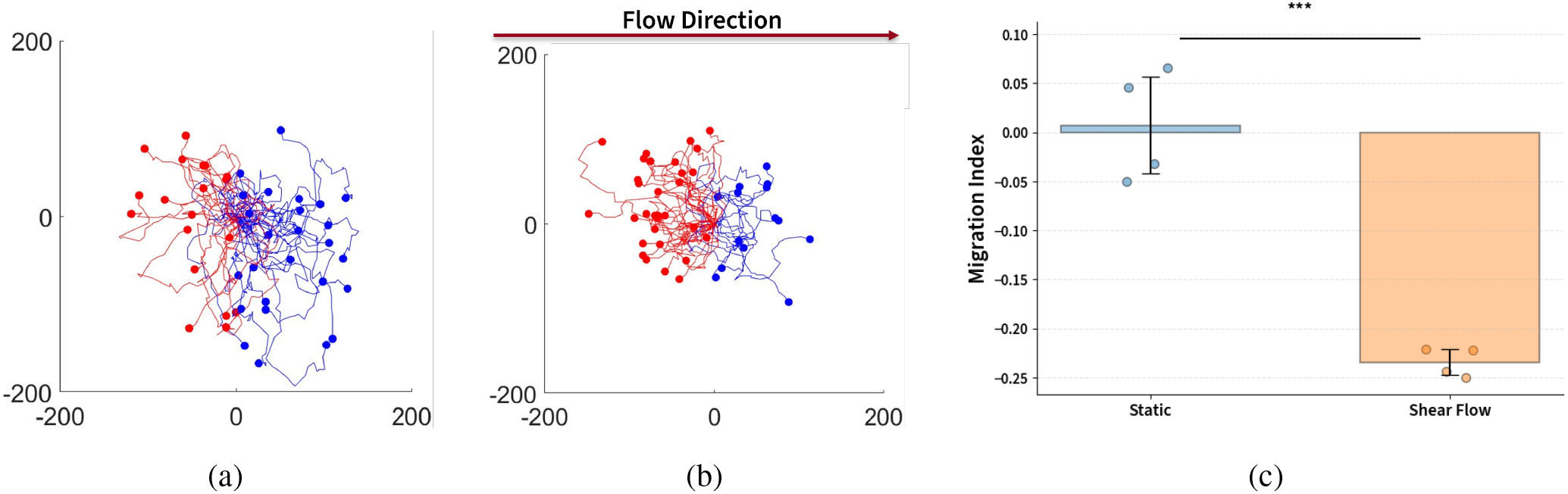
Migration behavior of KG1a cells on ICAM-1–coated polyacrylamide hydrogels under static and shear flow conditions. (A) Trajectories of KG1a cells under static conditions. (B) Trajectories of KG1a cells exposed to 400 s^−1^ shear flow. Arrow on the plot shows the direction of the flow. (C) Migration indices of cells under static and shear flow conditions. Cells exposed to shear flow exhibit significant upstream migration relative to random migration observed under static conditions (^***^p < 0.001).

In contrast, when subjected to a shear flow of 400 s^−1^, KG1a cells demonstrated pronounced upstream migration against the direction of flow. Time-lapse imaging and trajectory mapping revealed a pronounced directional bias, with most cells moving upstream over the course of the experiment (Figure 2b). Quantitative analysis of the MI under shear flow showed a significantly negative value of -0.234 ± 0.012 (Figure 2c), which is indicative of migration against the direction of flow. A corresponding video of upstream migration is available in the supplement (Video S2). Statistical comparison between the static and shear flow conditions using an unpaired t-test confirmed that this difference was highly significant (p < 0.001).

Together, these results validate that ICAM-1 functionalized polyacrylamide hydrogels can reliably support upstream migration in KG1a cells under shear flow. This system provides a robust platform for investigating migration behavior under physiologically relevant flow conditions.

### KG1a cells generate heterogenenous traction forces under static conditions

To establish a baseline for interpreting force generation during upstream migration, we first quantified the spatial and temporal characteristics of traction forces exerted by KG1a cells migrating on ICAM-1 functionalized hydrogels in the absence of flow. We imaged individual cells over a 30-minute period and extracted bead displacement fields using fluorescent microscopy and LIBTRC. Figures 3a – 3c show representative traction maps taken at 5-minute intervals for three distinct cells. Higher magnification images of each of the seven images for 3b are shown in the supplement, figure S1-S7. Force vectors and corresponding pseudocolor stress maps are overlaid onto cell footprints. While some cells exhibited strong focal forces at isolated regions, others displayed more diffuse or fragmented patterns of force engagement across the cell body. This variability serves as an important reference for evaluating how traction behaviors may reorganize under directional cues.

**Figure 3:**
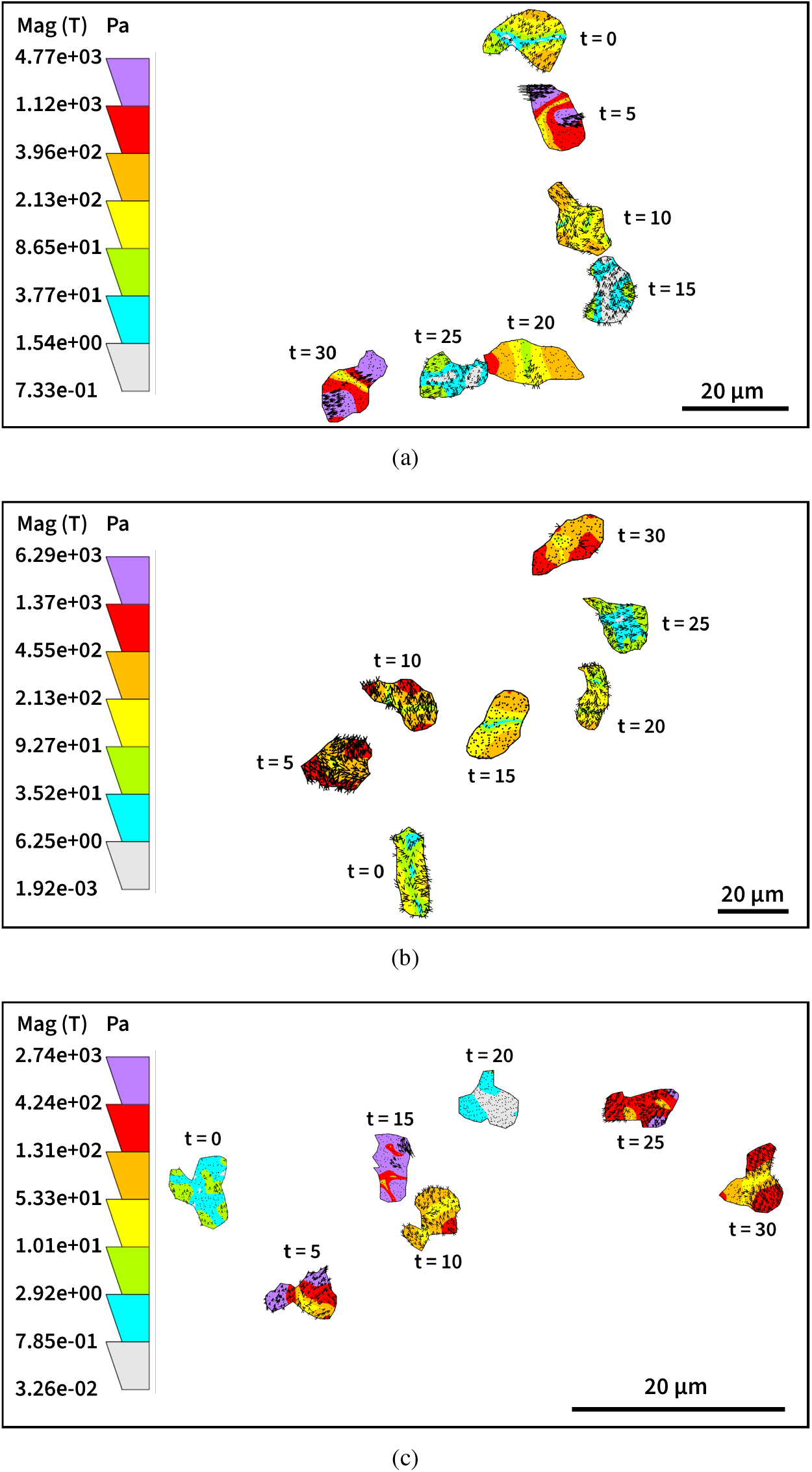
Traction force maps of KG1a cells under static conditions. Representative TFM images from three individual cells migrating on ICAM-1-functionalized hydrogels without flow. Images are shown at multiple time points over a 30-minute period. Force vectors (black arrows) and stress magnitudes (color overlays) highlight heterogeneity in spatial localization and intensity of traction forces.

To quantify the time-dependent dynamics of force generation under static conditions, we calculated the RMS traction force for each cell at 5-minute intervals over the 30-minute imaging period. Figure 4 shows the resulting RMS force profiles for the same three cells depicted in the TFM maps. Across cells, we observed substantial variability in both magnitude and temporal distribution. While one cell maintained relatively steady force output, the others exhibited pronounced oscillations, with RMS values rising and falling in roughly alternating cycles. Notably, cells 1 and 3 showed multiple transient force peaks spaced approximately 10-15 minutes apart, suggesting rhythmic engagement and release of traction forces. This pattern is consistent with a form of cyclic motility, potentially reflecting internal persistence times associated with protrusion-retraction cycles.

**Figure 4:**
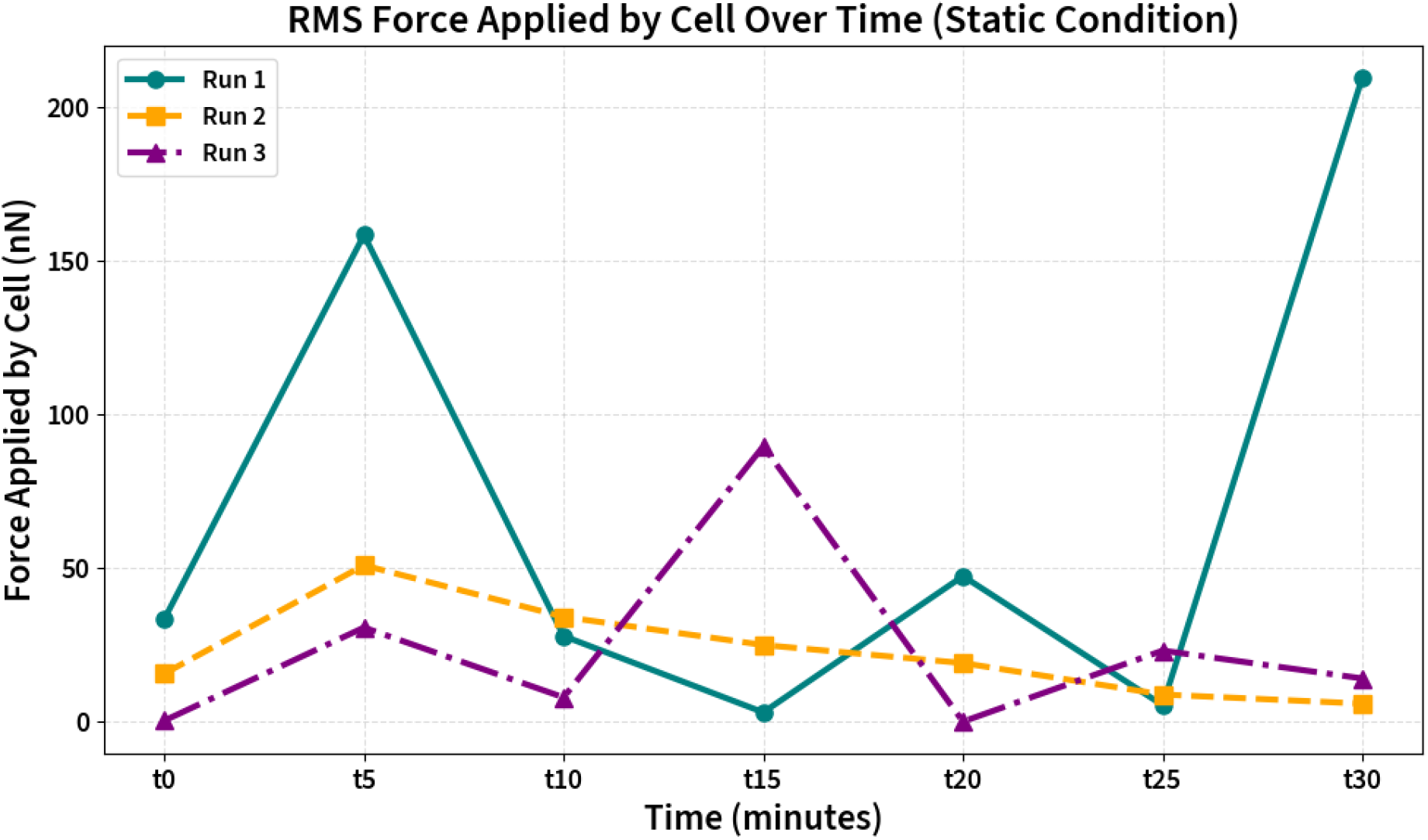
RMS force dynamics of KG1a cells under static conditions. Root-mean-square (RMS) traction force was computed at 5-minute intervals over a 30-minute window for three representative cells migrating without flow. Force magnitudes show fluctuating trends with no consistent polarization or temporal coordination.

Despite their differences, all cells generated measurable forces and engaged the substrate in a time-varying manner. However, in the absence of direction cues such as flow, these forces lacked spatial coherence or directional persistence. The distribution of traction remained heterogeneous across both the cell body and time course, hinting at a model in which KG1a cells operate in a mechanically active but directionless “search mode” under static conditions. In this regime, force generation is dynamic and locally intense but globally uncoordinated, providing a flexible baseline from which more structured migration behaviors may emerge under flow.

### KG1a cells exhibit temporally structured traction force dynamics during upstream migration

To examine how shear flow influences force generation in migrating leukocytes, we analyzed traction force dynamics in KG1a cells undergoing migration on ICAM-1 functionalized polyacrylamide gels under a shear rate of 400 s^−1^. Time-lapse TFM imaging was performed over a 30-minute window, with traction forces calculated at 5-minute intervals. Figures 5a – 5d display representative stress maps from three individual upstream-migrating cells. Higher magnification images of each of the six images for 5b are shown in the supplement, figure S8-S13. In all examples, traction forces were spatially organized along the axis of movement, with stress concentrated at discrete regions of the cell body. Elevated stress concentrations frequently appeared near the anterior of the cell, consistent with lamellipodial protrusions, while the rear – corresponding to the uropodia – exhibited comparatively reduced force. This spatial polarization contrasts with the fragmented and fluctuating force distributions observed under static conditions and suggests that upstream migration involves persistent, directional substrate engagement. While the precise morphology varied between cells, the emergence of pronounced, stable high-force zones at the leading edge indicates a fundamental reorganization of how traction is applied under flow.

**Figure 5:**
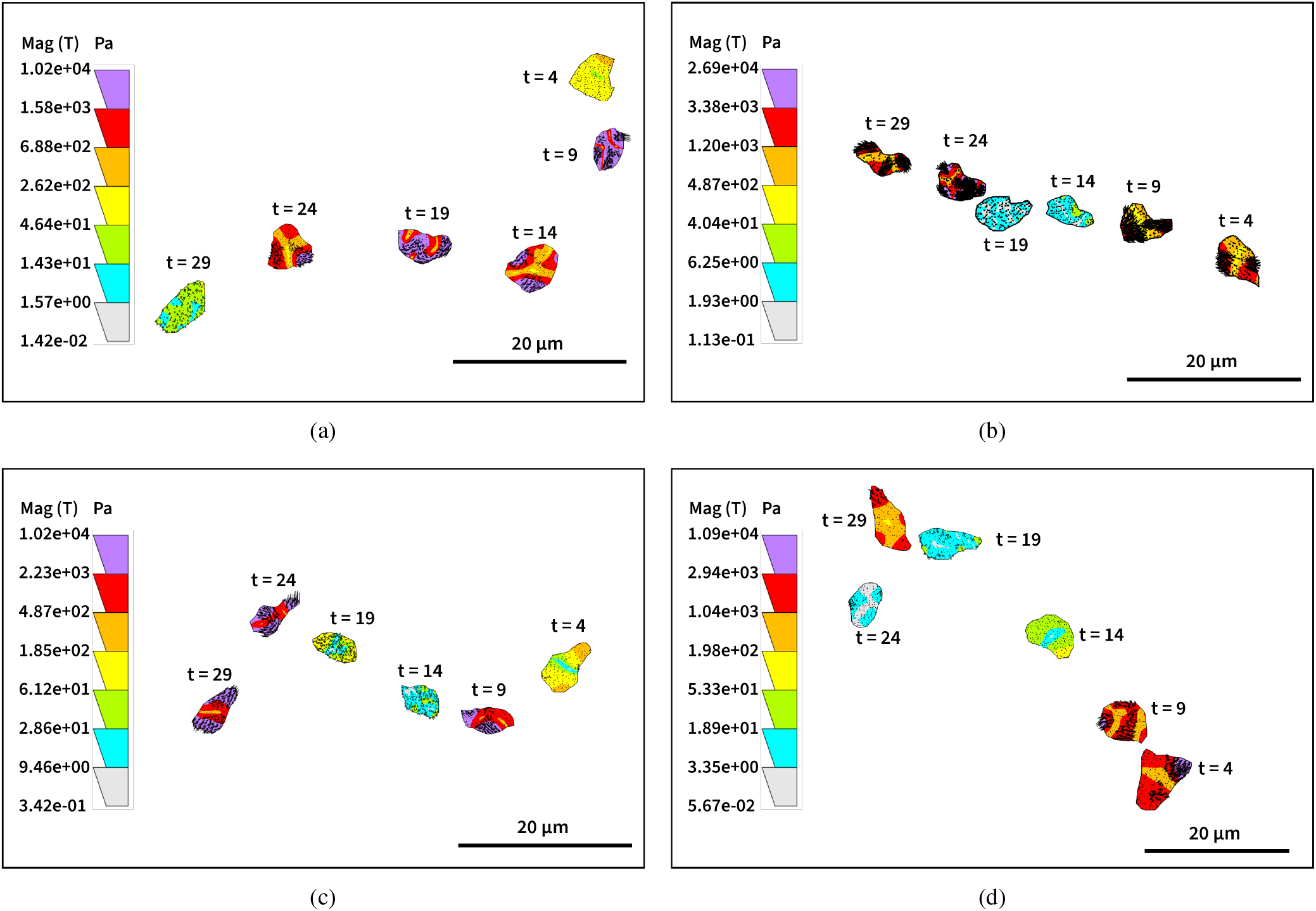
Traction force maps of KG1a cells during upstream migration. Representative TFM images from four KG1a cells migrating against shear flow on ICAM-1-coated hydrogels. Images were acquired at 5-minute intervals over a 30-minute time course. Stress magnitudes (color overlays) and traction vectors (black arrows) reveal distinct, directionally aligned zones of force generation. Compared to static migration, upstream cells exhibit stronger front-rear polarization and more persistent tractions over time.

To assess how traction force magnitude evolves over time in this context, we computed the RMS traction force at each 5-minute interval for the same three cells shown in Figures 5a – 5d. The resulting RMS force profiles are shown in Figure 6. Rather than maintaining a steady level of force, cells exhibited pulsatile force dynamics characterized by transient peaks separated by periods of reduced output, although the regularity and magnitude of these surges varied between individual cells. These surges were not randomly distributed; rather, they often corresponded to noticeable remodeling events in traction maps, such as the emergence of a new high-force zone or consolidation of existing traction at the leading edge. This temporal coupling between peak force magnitude and spatial redistribution supports a model in which upstream migration proceeds through cyclic episodes of traction buildup and release. Though not strictly periodic, the recurrence of these pulses may reflect underlying contractile cycles intrinsic to upstream motility.

**Figure 6:**
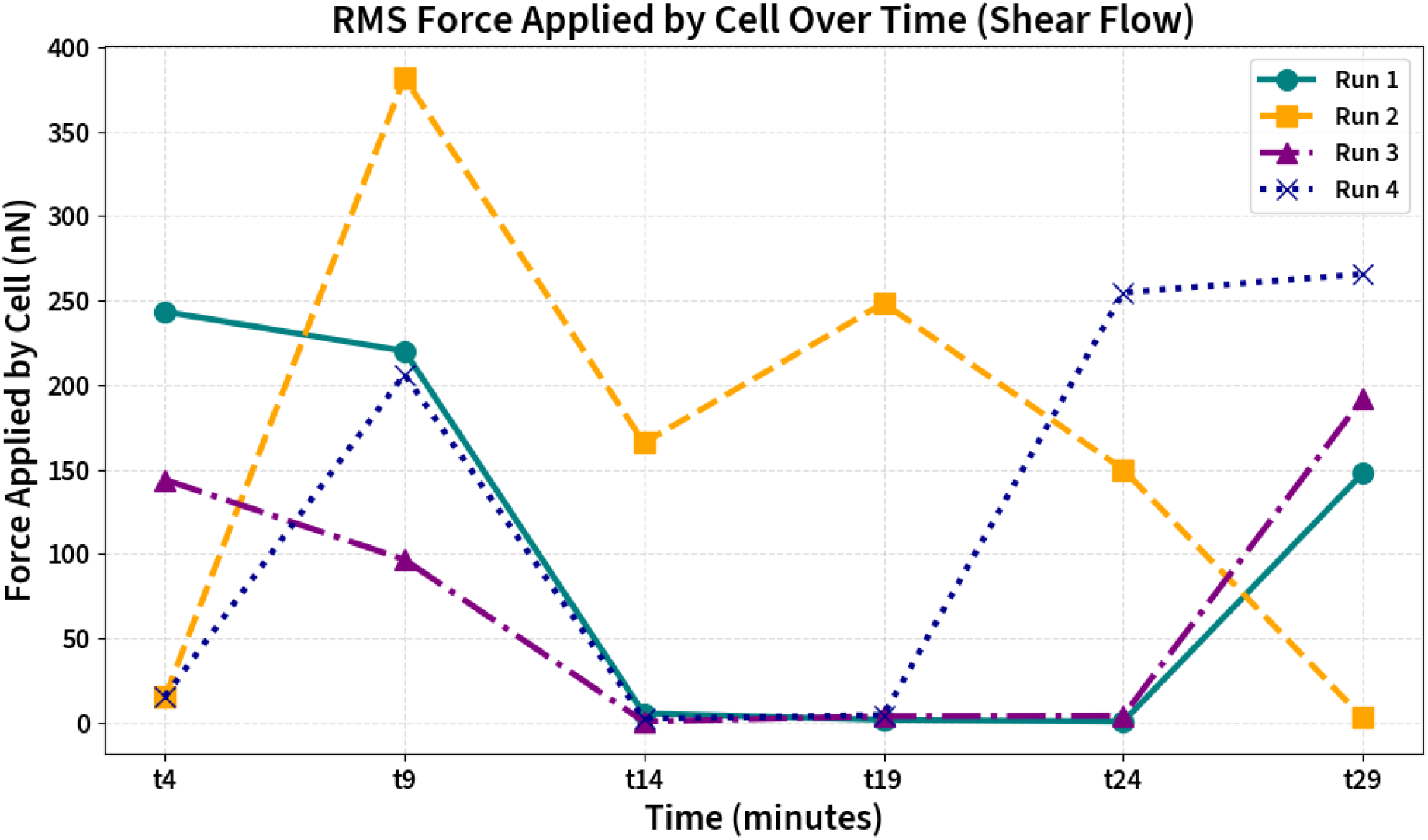
RMS force dynamics of KG1a cells during upstream migration. Root-mean-square (RMS) traction force values were calculated at 5-minute intervals for three individual KG1a cells migrating under flow. Traces reveal temporally structured force dynamics, with some cells exhibiting sharp transient peaks and others displaying gradual modulations. These fluctuations often correspond to morphological changes in the traction maps and reflect the mechanical tuning required to maintain directional migration against fluid shear.

While cyclic force modulation was also observed in some static cells, the dynamics exhibited during upstream migration were distinct in both organization and alignment. Under static conditions, force surges appeared more sporadic and spatially uncoordinated, often shifting unpredictably across the cell body. In contrast, force peaks during upstream migration tended to coincide with sustained traction at the leading edge, aligning with the axis of movement and suggesting a more coherent integration of contractile activity with migration direction. This distinction implies that while intrinsic contractile cycles may be a shared feature of KG1a motility, exposure to flow may transform these cycles from a randomized exploratory behavior into a mechanically reinforced, directionally biased migration.

The emergence of both spatial coherence and temporal modulation during upstream migration points to a shift in how traction forces are coordinated. Rather than being governed by internal cues alone, the presence of flow appears to shape the organization of mechanical output, biasing force generation toward specific regions of the cell and tuning its temporal structure. These features may reflect an adaptation that enables leukocytes to overcome fluid shear and persist in upstream migration. This sets the stage for a broader population-level analysis to determine whether the observed trends – greater amplitude, directionality, and temporal structure – are conserved across migrating cells.

### Upstream migration is associated with increased traction force magnitude at the population level

The temporal and spatial analyses of individual KG1a cells suggest that upstream migration is accompanied by reorganized and intensified traction force dynamics. To test whether these trends extend beyond isolated examples, we performed a population-level comparison of RMS traction forces across static and upstream conditions. For each cell, we extracted minimum, maximum, and average RMS values over the 30-minute imaging period. These metrics capture distinct aspects of force behavior – baseline adhesion strength, peak contractile output, and sustained force levels, respectively – and offer a comprehensive view of how traction generation differs between migratory modes.

Figure 7 summarizes the distribution of force magnitudes across conditions, with n=12 cells analyzed for upstream migration and n=11 cells for static condition. Cells migrating upstream exhibited significantly greater maximum RMS forces than those under static conditions (mean: 428.5 ± 63.0 nN vs 220.8 ± 22.2 nN, p = 0.0078), indicating that upstream migration imposes greater contractile demands on the cell. Similarly, the average RMS force per cell was elevated in the upstream group (mean: 82.6 ± 12.9 nN vs 45.9 ± 4.4 nN, p = 0.0184), reflecting a shift toward more sustained mechanical engagement with the substrate. In contrast, minimum RMS force values were statistically indistinguishable between the two conditions (p = 0.2600), suggesting that the fundamental substrate engagement remains unchanged, and the key differentiator lies in the peak and average force output.

**Figure 7:**
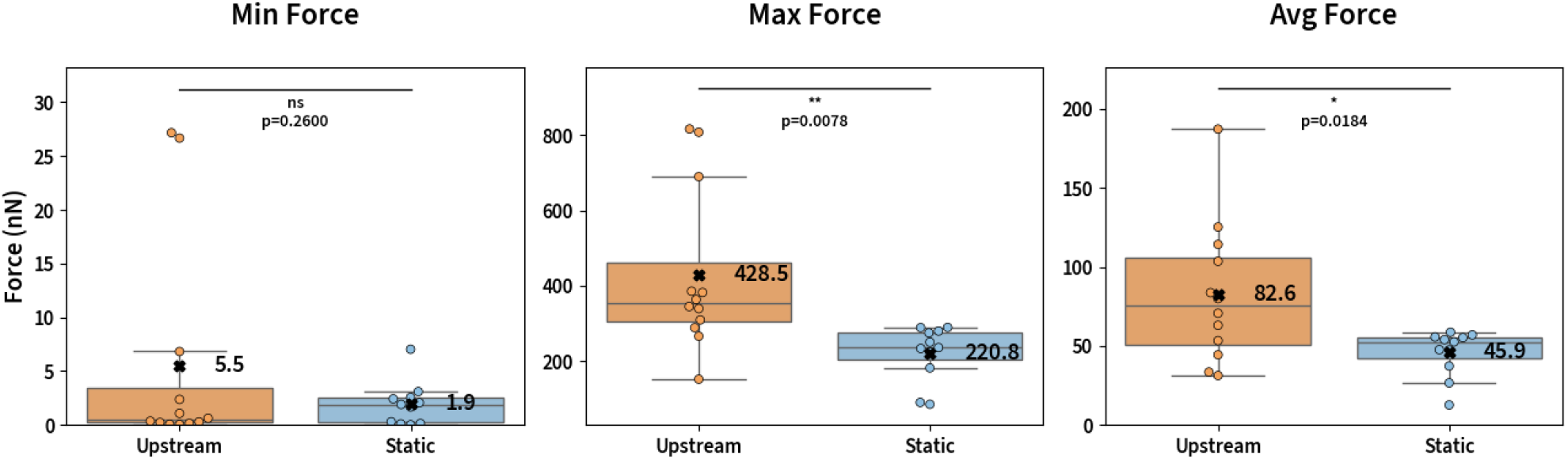
Population-level comparison of traction force metrics between upstream and static migration conditions. Root-mean-square (RMS) traction force values were quantified across all analyzed cells for both conditions (upstream: n=12; static: n=11). Box-and-whisker plots show distributions of minimum, maximum, and average RMS forces per cell. While minimum forces were comparable between groups (*p* = 0.2600, *ns*), upstream-migrating cells exhibited significantly higher maximum forces (*p* = 0.0078, **), and moderately increased average forces (*p* = 0.0184, *). These population-level trends support the notion that upstream migration involves greater mechanical output relative to random motility under static conditions.

These findings affirm that the mechanical demands of upstream migration are not only qualitatively distinct, but also quantitatively enhanced at the population level. The elevation in maximum and average traction forces is consistent with the need to resist external shear while maintaining directional motility. This increased force generation likely reflects both intensified cytoskeletal contraction and more stable integrin engagement at the leading edge, consistent with the single-cell patterns observed in our time-lapse analyses. Furthermore, the preservation of baseline force levels suggests that upstream migration does not merely involve “turning up” global adhesion but rather tunes specific components of the force program – enhancing output selectively without disrupting fundamental substrate contact.

Taken together, these results demonstrate that upstream migration represents a mechanically reinforced mode of cell motility, characterized by amplified and spatiotemporally coordinated traction force production. This population-level shift supports the interpretation that shear flow is not simply tolerated but actively integrated into the mechanical program of leukocyte migration, transforming a stochastic search pattern into a directed and force-intensive migration behavior.

## DISCUSSION

In this study, we combined time-resolved traction force microscopy with flow-adapted hydrogel substrates to examine the mechanics of upstream migration in KG1a cells. By comparing cells migrating under static conditions to those exposed to physiologically relevant shear flow, we identified distinct patterns of force generation across space and time. Our findings suggest that upstream migration is not simply a reversal of directionality, but a reprogramming of the cell’s traction machinery into a more amplified, polarized, and temporally structured force regime. These observations highlight a mechanically reinforced migratory state that enables leukocytes to sustain movement against fluid shear.

At the single-cell level, upstream-migrating KG1a cells exhibited traction forces that were both spatially and temporally organized. Unlike static conditions, where tractions were fragmented and shifted rather unpredictably, upstream migration was characterized by persistent high-force zones localized to the leading edge. This spatial coherence was accompanied by rhythmic force surges – transient RMS force peaks that often coincided with traction reorganization – suggesting a cycle of mechanical buildup and release that may support continued propulsion under flow. While some cells in static conditions also exhibited cyclic force patterns, these were more sporadic and lacked consistent spatial alignment, reinforcing the idea that external flow transforms a pre-existing exploratory behavior into a directed, force-intensive migratory program.

At the population level, upstream migration was associated with a marked increase in both average and maximum traction force magnitudes, while minimum force values remained similar between the two populations. These shifts suggest that upstream migrating cells amplify their mechanical engagement specifically during active phases of migration, rather than globally elevating substrate engagement. The conservation of baseline force levels implies that cells preserve low-force states but punctuate them with intensified force bursts when migration demands it. This selective tuning of mechanical output remains consistent with our single-cell findings and underscores the adaptability of the migratory machinery in response to external constraints.

These mechanical adaptations likely serve a critical biological function. Leukocytes that engage in upstream migration – such as T cells (9), hematopoietic progenitors (14), or neutrophils when Mac-1 is blocked (15) – must navigate shear environments within postcapillary venules. The ability to polarize force output and elevate contractility at the leading edge may allow these cells to maintain adhesion while overcoming drag, particularly in the absence of chemotactic gradients. This may represent a specialized form of mechanotaxis – where force, rather than ligand distribution, drives directional persistence. Supporting this, Valignat and coworkers proposed the “wind vane” model in which the non-adherent uropod of T cells acts as a passive sensor, aligning the cell’s polarity with the direction of fluid flow, a morphology that was directly observed by confocal imaging in upstream-migrating T-cells (26). Importantly, Valignat and coworkers also demonstrated that the stress generated by the cell during migration (on the order of 1 nN, driven by actin polymerization) far exceeds the hydrodynamic force imposed by physiological shear flow, which is estimated at only 0.4 - 0.6 nN (26). In our experiments, a similar disparity is observed: the hydrodynamic drag on a KG1a cell is approximately 0.4 nN under our flow conditions, whereas the traction forces we measured during upstream migration averaged at 82.6 ± 12.9 nN, orders of magnitude above the external load. Taken together, these findings support the notion that upstream migration is not simply a matter of resisting flow, but is instead enabled by the cell’s intrinsic capacity to generate and sustain contractile forces, well beyond the levels imposed by the surrounding flow. While we did not measure traction stresses in T cells directly, we plan to compare traction stresses among cell types to better understand how the mechanical programs operate during upstream migration of different cell types.

It might be informational to compare the magnitudes of RMS forces observed here to those previously reported for migrating neutrophils and macrophages (19, 20). Under a gradient of fMLP of 10 µM/10 µM, neutrophils displayed an RMS force of ∼100 nN, while macrophages undergoing random motility displayed an RMS force of ∼1200 nN. The forces we observe here for directional motion of KG1a cells against flow fall between these values, and most notably exceed those values reported chemotactically migrating neutrophils.

In addition to its biological findings, this study advances the technical toolkit for studying leukocyte migration in mechanically active environments. By capturing time-resolved RMS force traces, our approach enables the identification of dynamic features such as force surges, temporal modulation, and front-rear polarization, offering a richer phenotypic readout than static force maps alone. The method provides an interpretable and quantitative framework well-suited for small-scale perturbation studies. This force-based readout could be particularly valuable in focused CRISPR or pharmacological screens targeting cytoskeletal or adhesion regulators, where mechanical output itself serves as a functional phenotype.

Looking forward, we see several promising extensions of this work. One key opportunity lies in identifying the molecular determinants of force modulation during upstream migration. Our platform offers a path to mechanistic dissection by linking mechanical phenotypes to specific molecular regulators. Recent work by Roy and coworkers has demonstrated that both the Crk adaptor proteins and the E3 ubiquitin ligase c-Cbl are critical for upstream migration in T cells. Loss of Crk proteins disrupts LFA-1-dependent actin polymerization, cell spreading, and mechanosensing, abrogating the ability of T cells to migrate upstream on ICAM-1 (16). Deletion of c-Cbl, meanwhile, selectively impairs upstream migration without altering cell morphology, suggesting that c-Cbl acts through a distinct signaling mechanism downstream of LFA-1. Several additional candidate regulators – such as those involved in integrin recycling, mTOR signaling, and actin dynamics – have also emerged from recent genome-wide CRISPRi screens in neutrophil-like HL-60 cells by the Theriot lab (27). Leveraging our traction force framework to assay these regulators could illuminate force-based modes of migration control and clarify the transition between undirected and directed migration programs.

In summary, this study provides a quantitative foundation for understanding how leukocytes adapt their mechanical behavior to migrate upstream under flow. By integrating spatial, temporal, and population-level analyses of traction force, we show that upstream migration is a distinct biophysical state – not just in direction but in how cells generate and coordinate force. This work lays the foundation for future studies uncovering the molecular programs that drive force adaptation during upstream migration, and sets the stage for force-based phenotyping as a central readout in targeted screens for migratory immune cell function.

## CONCLUSION

This study demonstrates that upstream migration in leukocytes involves a distinct mode of mechanical engagement, characterized by spatially polarized and temporally modulated traction forces. Using time-resolved TFM under physiologically relevant shear flow, we show that KG1a cells reorganize their traction dynamics into a force-amplified, directionally coherent state that enables sustained migration against flow. These mechanical signatures differ markedly from the more stochastic and fragmented behaviors observed under static conditions. By quantifying single-cell and population-level force patterns, we provide a framework for interpreting upstream migration as a biophysically reinforced state. This platform sets the stage for identifying the molecular regulators that coordinate mechanical output with migration under flow, and offers a force-resolved readout for targeted mechanistic perturbations in immune cells.

## AUTHOR CONTRIBUTIONS

Dong-hun Lee performed the experiments and the data analysis. Daniel A. Hammer designed the research. Dong-hun Lee and Daniel A. Hammer wrote the article.

## ACKNOWLEDGMENTS

This work was funded by NIH R01 grant GM143357. We thank Micah Dembo for graciously sharing the LIBTRC software used for this research.

## SUPPLEMENTARY MATERIAL

An online supplement to this article can be found by visiting BJ Online at http://www.biophysj.org.

